# The effects of age on neural correlates of recollection: transient versus sustained fMRI effects

**DOI:** 10.1101/2023.04.12.536508

**Authors:** Mingzhu Hou, Marianne de Chastelaine, Michael D. Rugg

## Abstract

Prior fMRI findings in young adults indicate that recollection-sensitive neural regions dissociate according to the time courses of their respective recollection effects. Here, we examined whether such dissociations are also evident in older adults. Young and older participants encoded a series of word-object image pairs, judging which of the denoted objects was the smaller. At test, participants first judged whether a test word was old or new. For items judged old, they were required to recall the associated image and hold it in mind across a variable delay period. A post-delay cue denoted which of three judgments should be made on the retrieved image. Older adults demonstrated significantly lower associative memory performance than young adults. Replicating prior findings, transient recollection effects were identified in the left hippocampus, medial prefrontal cortex and posterior cingulate, while sustained effects were widespread across left lateral cortex and were also evident in the bilateral striatum. With the exception of those in the left insula, all effects were age-invariant. These findings add to the evidence that recollection-related BOLD effects in different neural regions can be temporally dissociated. Additionally, the findings suggest that both transient and sustained recollection effects are largely stable across much of the healthy adult lifespan.

## 1. Introduction

Compared with young adults, older adults typically demonstrate reduced episodic memory performance – that is, a decline in the ability to recollect unique events (Nilsson, 2003; Nyberg and Pudas, 2019). This well documented finding has motivated numerous functional magnetic resonance imaging (fMRI) studies that examined the effects of age on the neural correlates of episodic memory encoding and retrieval (for review, see Maillet and Rajah, 2014; Tromp et al., 2015; Wang and Cabeza, 2016). Here we focus on fMRI ‘recollection effects’, the enhanced neural activity elicited by retrieval cues giving rise to successful as opposed to failed recollection. We describe what, to our knowledge, is the first fMRI study to contrast the temporal properties of recollection effects in young and older adults.

fMRI recollection effects have been extensively examined in a wide variety of experimental paradigms (for reviews, see Kim 2010, 2013; Rugg and Vilberg, 2013; Rugg et al., 2015). Successful recollection has consistently been associated with enhanced BOLD activity in several cortical regions (collectively termed the ‘core recollection’ (Rugg and Vilberg, 2013) or ‘posterior medial’ (Ranganath and Ritchey, 2012) networks) including the medial prefrontal cortex (mPFC), posterior cingulate (PCC)/retrosplenial cortex, hippocampus, parahippocampal gyrus (PHC), left middle temporal gyrus (MTG) and left angular gyrus (AG).

Of importance, with only two exceptions, fMRI studies examining recollection effects have employed memory tests in which mnemonic information could be used to guide behavior as soon as the information had been retrieved. Such tests provide no opportunity to contrast the temporal properties of the recollection effects manifest in different brain regions. In the two exceptions (Vilberg and Rugg, 2012, 2014), young participants were required to maintain recollected information over a variable delay interval that was terminated by a cue that signaled which of three judgments should be made about the recollected study episode (this procedure was adopted to encourage participants to maintain a representation of the studied item across the interval). The authors reported that the hippocampus, PHC, mPFC and PCC demonstrated ‘transient’ recollection effects that were independent of the duration of the delay interval. By contrast, the striatum, left inferior frontal gyrus (IFG), inferior temporal gyrus (ITG), intraparietal sulcus (IPS) and, among the members of the core recollection network, the left AG and MTG demonstrated ‘sustained’ effects that tracked the delay interval.

Prior studies comparing recollection effects in cognitively healthy young and older adults have reported largely age-invariant effects in core recollection regions (de Chastelaine et al., 2016; Dulas and Duarte, 2012; Folville et al., 2020; Hou et al., 2021, 2022; Wang et al., 2016), especially when memory performance was matched or statistically equated between age groups (for exceptions, see Angel et al., 2013; Daselaar et al., 2006). These findings suggest that recollection effects are largely unaffected by increasing age, but the time courses of the effects have yet to be examined in older adults. Thus, it is unknown whether recollection effects remain age-invariant when the retrieved information must be maintained over an interval, and hence require an interaction between episodic and working memory, before it is used to guide a memory-based decision.

Here, we employed an experimental procedure closely similar to that of Vilberg and Rugg (2012, 2014) with the aim of examining the effects of age on transient and sustained recollection effects. We expected to replicate prior findings of regional dissociations in the time-courses of recollection effects in young adults. Given prior findings of null or minimal age differences in the recollection effects elicited in conventional memory tests (see above), we predicted that transient recollection effects would be largely age-invariant. At issue is whether this null finding extends to sustained effects. Notably, evidence of an age-related attenuation of sustained recollection effects would significantly qualify prior findings that the neural correlates of recollection are age-invariant and, in addition, identify regions whose mnemonic functions might be vulnerable to increasing age.

## 2. Materials and methods

### 2.1 Participants

Participants were 24 young (18-29 years old) and 24 older (65-75 years old) adults recruited from the UT Dallas and surrounding metropolitan Dallas communities. They were right-handed fluent English speakers, had normal or corrected-to-normal vision, no history of neurological or psychiatric illness and were not taking prescription medications that affected the central nervous system. Informed consent was obtained in accordance with UT Dallas Institutional Review Board guidelines. Participants were compensated at the rate of $30 an hour. Data from 3 participants (1 young, 2 older) were excluded from all analyses due to insufficient trial numbers (< 5) for one or more events of interest (see below).

### 2.2 Neuropsychological testing

A standardized neuropsychological battery was administered on a separate day from the MRI session. The test battery comprised the Mini-Mental State Examination (MMSE), the California Verbal Learning Test-II (CVLT; Delis et al., 2000), Wechsler Logical Memory (Wechsler, 2009), the Symbol Digit Modalities test (SDMT; Smith, 1982), the Trail Making Test A and B (Reitan and Wolfson, 1985), the Digit span forward and backward test (Wechsler, 1981), the F-A-S letter fluency test (FAS; Spreen and Benton, 1977), the Category Fluency test (Benton, 1968) and Raven’s Progressive Matrices (List 1; Raven et al., 2000). Forty-six participants completed the Wechsler Test of Adult Reading (WTAR, Wechsler, 2001) and 2 others completed its revised version, the Wechsler Test of Premorbid Functioning (TOPF; Wechsler, 2011). The TOPF and WTAR scores were converted to commensurate measures by scaling the raw scores according to the two test’s respective age-scaled norms for age 45 years (the mean age of our participants). To reduce the likelihood of including participants with mild cognitive impairment, potential participants were excluded from the MRI session if they had a score on the MMSE < 26, a score on a standardized memory test >1.5 SD below the age-appropriate norm, or low performance (>1.5 SD below norm) on two or more of the non-memory tests.

### 2.3 Experimental items and procedure

An associative memory task was undertaken in the MRI scanner. The task comprised 4 scanned study-test blocks. Separate stimulus lists were created for yoked pairs of young and older participants. Items for the study phase were 80 word-object image pairs randomly selected from a pool of 120 such pairs. The test items comprised all 120 words from the pool. Two fillers were presented at the beginning of each study and test block. An additional 48 words and 32 images were used as practice items.

Figure 1 illustrates a schematic depiction of the associative memory task. The task requirement at study was to imagine the objects denoted by the word and the image interacting with each other in real life, and to signal which object was the smaller by pressing one of two keys. Each word-image pair was presented for 5s and was preceded by a red fixation cross for 0.5s. A white fixation cross followed for 1.5s.

**Figure 1.**
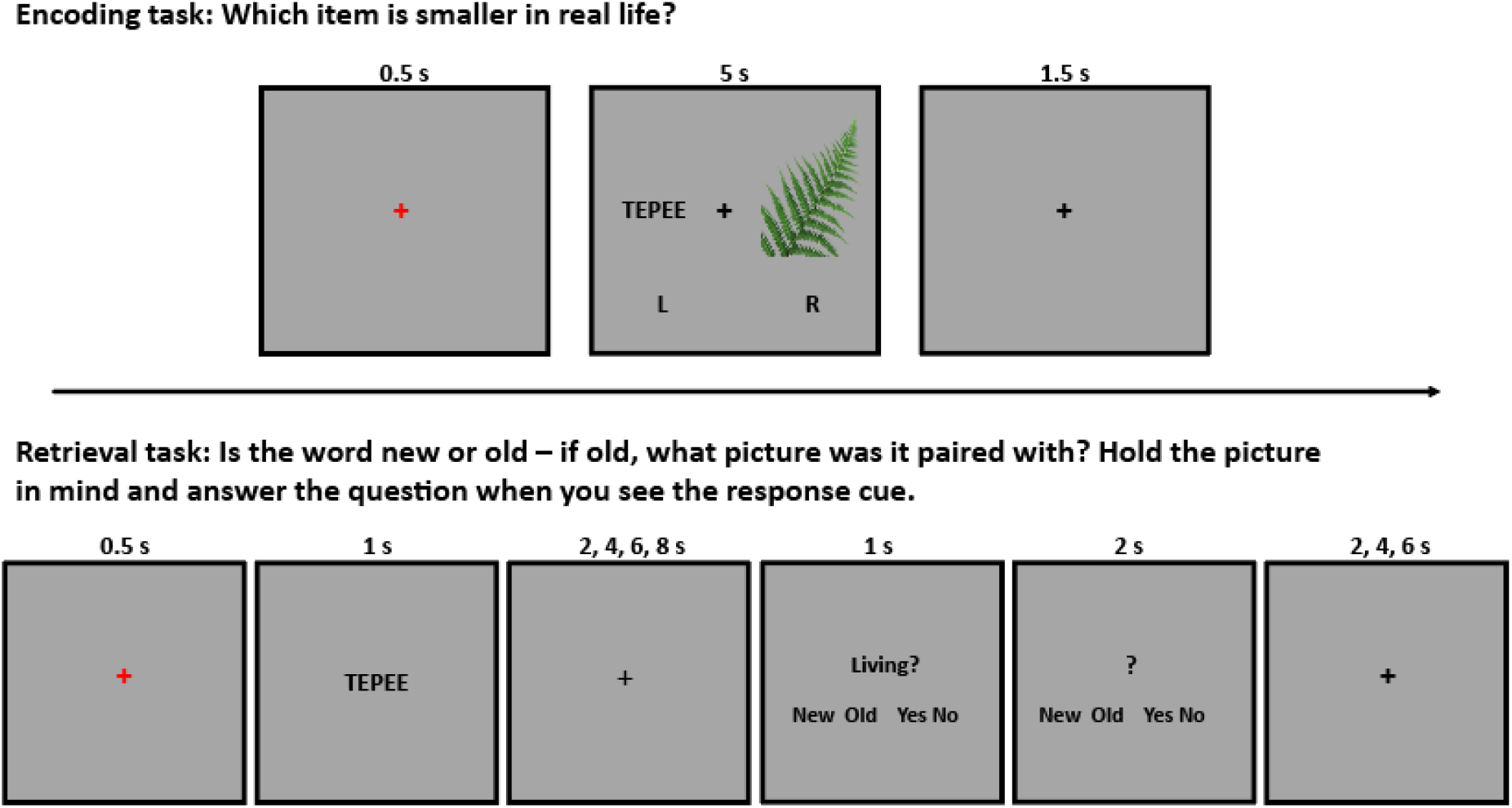
Schematic depictions of the study and test tasks.

Each of the 4 test blocks comprised 20 studied and 10 unstudied words. Instructions were to judge whether the word was old or new, and if old, to recall and hold in mind the image that had been paired with the word at study. After a variable delay period, a cue was presented that required one of three different judgments about the recalled object. The cues were ‘shoebox’, ‘living’, and ‘house’, respectively, signaling: Would the recalled object fit inside a shoebox? Is the recalled object living or part of a living thing? Would the recalled object be found inside a house? Participants were instructed to withhold their response until the appearance of the cue. They were required to press one of four keys that respectively signaled a ‘New’, ‘Old’, ‘Yes’ or ‘No’ response. Participants were to press ‘New’ if the test word was judged as new, ‘Old’ if the test word was judged old but the paired study image could not be recalled, or ‘Yes’ or ‘No’ to indicate their answer to the question posed by the cue about the recalled object. Responses were made using the index and middle fingers of both hands. The timing of the events comprising a test trial is given in Figure 1. The ordering of the test words was pseudo-randomized so that no more than 3 words from the same old/new category occurred in succession. The ordering of the cues was also pseudo-randomized so that the same question did not appear on more than three consecutive trials. For both study and test phases, response hand assignment was counterbalanced across participants. To ensure that participants understood the task requirements, practice on the experimental tasks was undertaken before they entered the scanner.

### 2.4 MRI acquisition and preprocessing

MRI data were acquired with a Siemens PRISMA 3T MR scanner equipped with a 32-channel head coil. Functional scans were acquired with a T2*-weighted echoplanar (EP) sequence (TR 1520 ms, TE 30 ms, flip angle 70°, field-of-view (FOV) 220 × 220, matrix size 80 × 80, multiband factor = 2). Each EP volume comprised 50 slices (2.5 mm thickness, 0.5 mm inter-slice gap) with an in-plane resolution of 2.5 × 2.5 mm and was acquired with an anterior-to-posterior phase encoding direction. A T1-weighted image was acquired with an MP-RAGE pulse sequence (TR = 2300 ms, TE = 2.26 ms, FOV = 256 x 256, voxel size = 1 x 1 x 1 mm, 160 slices, 0.5mm inter-slice gap, sagittal acquisition). To allow correction for geometric distortions in the EP images caused by magnetic field inhomogeneities, a field map was acquired after the functional scans using a double-echo gradient echo sequence (TE 1/2 = 4.92 ms/7.38 ms, TR = 520 ms, flip angle = 60°, FOV 220 × 220, 50 slices, 2.5mm slice thickness). Functional images were acquired at both study and test, but here we only report findings from the test phase.

Data preprocessing was performed with SPM12 implemented in MATLAB R2018b. The functional volumes were field-map corrected, realigned to the across-run mean image and spatially normalized to a study-specific template (see de Chastelaine et al., 2015, for a description of template construction). The normalized images were resampled to 2.5 mm isotropic voxels and smoothed with a 6 mm Gaussian kernel. Before being entered into subject-wise General Linear Models (GLMs), the functional data from the four test blocks were concatenated using the spm_concatenate.m function. Anatomical images were normalized to a study-specific T1 template following procedures analogous to those applied to the functional images.

### 2.5 Behavioral analyses

For the associative memory task, the proportions of associative hits (old words for which a correct answer was given in response to the cue), associative misses (old words for which an incorrect or an ‘old don’t recall’ answer was given in response to the cue) and correct rejections (new words correctly judged new) were subjected to a 3 (judgment type) x 2 (age group) ANOVA. For the reaction times (RTs) related to these responses, an analogous ANOVA (factors of judgment type, age group) was implemented. An additional 4 (delay interval: 2, 4, 6, 8s) x 2 (age group) ANOVA was conducted to examine associative hit rates as a function of delay interval.

We also computed Pr [p(associative hit) + p(associative miss) – p(false alarm)] as an estimate of item memory. An independent sample t-test was conducted to assess possible age differences in item memory.

Statistical analyses were conducted with SPSS 27.0. Non-sphericity between the levels of repeated measures factors in the ANOVAs was corrected with the Greenhouse–Geisser procedure (Greenhouse and Geisser, 1959). Significance levels for all tests were set at p < 0.05.

### 2.6 fMRI analyses

We employed a relatively liberal cutoff of 5 trials per event of interest (the results were however largely unchanged when the cutoff was raised to 10 trials; see supplementary materials). As noted previously, based on the 5-trial cutoff, data from 3 participants (1 young, 2 older) were excluded from the analyses.

To identify transient and sustained recollection effects, we constructed two sets of participant-level GLMs that, respectively, modeled the neural activity elicited by the test items with a delta function and a boxcar function (hereafter, delta-GLM and boxcar-GLM). The rationale for this approach was that the delta-GLM would be sensitive to transient activity, whereas the boxcar-GLM would capture activity that was sustained across the delay interval. We employed the two regressors in separate statistical models to eliminate any reduction in sensitivity due to collinearity between the regressors (the findings were, however, largely unchanged when we included the delta and boxcar functions in the same GLM).

For each participant-level GLM, three event types were modeled: associative hits, item hits (old words for which an ‘old don’t recall’ answer was given in response to the cue), and correct rejections. Other event types, together with fillers and trials with multiple or absent responses, were modeled as events of no interest. The models also included as covariates six regressors modeling motion-related variance (three for rigid-body translation and three for rotation) and four constants for means across test blocks. Data from volumes with a transient displacement (relative to the prior volume) of > 1 mm or > 1° in any direction were modeled as covariates of no interest.

For the delta-GLM, the delta functions onset concurrently with the onset of each test word and the resulting time series was convolved with SPM’s canonical hemodynamic response function (HRF). For the boxcar-GLM, the boxcar functions also onset concurrently with the onset of each test word, but varied between 3, 5, 7, and 9s in duration, corresponding to the duration of the test word (1s) and the subsequent delay interval (2, 4, 6 or 8s). As for the delta-GLM, the boxcar regressors were convolved with a canonical HRF.

For both GLMs, the participant-specific parameter estimates were taken forward to a second, across-participant, GLM. This took the form of a mixed effects ANOVA model, incorporating the factors of age group (young, older) and judgment type (associative hit, item hit, correct rejection). Effects that survived a height threshold of p < 0.001 and a FWE corrected cluster extent threshold of p < 0.05 (k > 59) were deemed significant. Given our *a priori* interest in the hippocampus (see Introduction), effects in this region were evaluated using a small volume correction within an anatomically defined bilateral hippocampal mask. In this case the cluster wise FWE corrected threshold corresponded to a cluster size of 10 or more voxels. Transient recollection effects were identified through a two-stage procedure. First, we identified clusters demonstrating associative hit > item hit effects from the delta-GLM. We then exclusively masked the outcome of this contrast with the corresponding contrast (height thresholded at p < 0.001) from the boxcar-GLM to remove any clusters where the recollection effects were sustained across the delay interval. Sustained recollection effects were identified by the associative hit > item hit contrast from the boxcar-GLM. Age-invariant transient and sustained recollection effects were identified by exclusively masking the across-group main effect with the corresponding age group x judgment type interaction contrast (conservatively thresholded at p < 0.05). Regions demonstrating age differences in transient or sustained effects were identified by the age group x judgment type interaction contrast (now thresholded as for the other whole brain contrasts; see above). For each cluster identified by the interaction contrast, we extracted the mean parameter estimates averaged across all voxels within a 3 mm (for hippocampus) or 5 mm (for neocortical regions) radius of the peak of the cluster. The parameter estimates were subjected to 2 (age group) x 2 (judgment type) mixed-design ANOVAs to elucidate the interactions.

In addition to the GLM approach described above (hereafter, the ‘standard’ GLM), we constructed an additional GLM in which a finite impulse response (FIR) model was employed to estimate the time courses of the BOLD responses elicited by the events of interest. The time courses were estimated across 14 TRs extending from 1 TR prior to test item onset to 13 TRs thereafter. Time courses associated with the 2, 4, 6, and 8s delay intervals were separately truncated at 5, 6, 7 and 8 TR post-item onset to avoid potential contamination with BOLD activity elicited on the following trial (cf. Vilberg and Rugg, 2012). The FIR model was used to visualize the BOLD time courses from the clusters identified by the standard GLMs, and to contrast the associative hit-related parameter estimates for the 2s and 8s delay intervals (see below). For transient recollection effects, we extracted mean parameter estimates from 5 mm radius spheres centered on the peak voxel of each of the neocortical clusters identified with the standard GLM. For the hippocampus, mean parameter estimates were derived from voxels falling within a 3 mm radius sphere centered on each peak. For the sustained effects, we selected the four exemplary left hemisphere regions (IFG, ITG, IPS and AG). For the first three of these regions, mean parameter estimates were extracted from a 5mm sphere centered on the peaks originally reported by Vilberg and Rugg (2012). For the AG, mean parameter estimates were extracted from an anatomically defined region of interest (ROI) comprising the Pga and Pgp subregions. The ROI was generated from the Anatomy toolbox v3.0 (Eickhoff et al., 2007). These regions are illustrated in Figure 4A and their coordinates are given in the figure legend. To examine whether recollection-related activity in the regions demonstrating sustained effects tracked the delay interval, we conducted a 2 (age group) x 2 (delay interval: 2s, 8s) x 11 (time point: 0-10 TRs after word onset) ANOVA on the parameter estimates for the associative hit trials derived from the FIR model. Significance levels for all tests were set at p < 0.05 and were family-wise corrected for multiple comparisons using the Bonferroni procedure.

**Figure 2.**
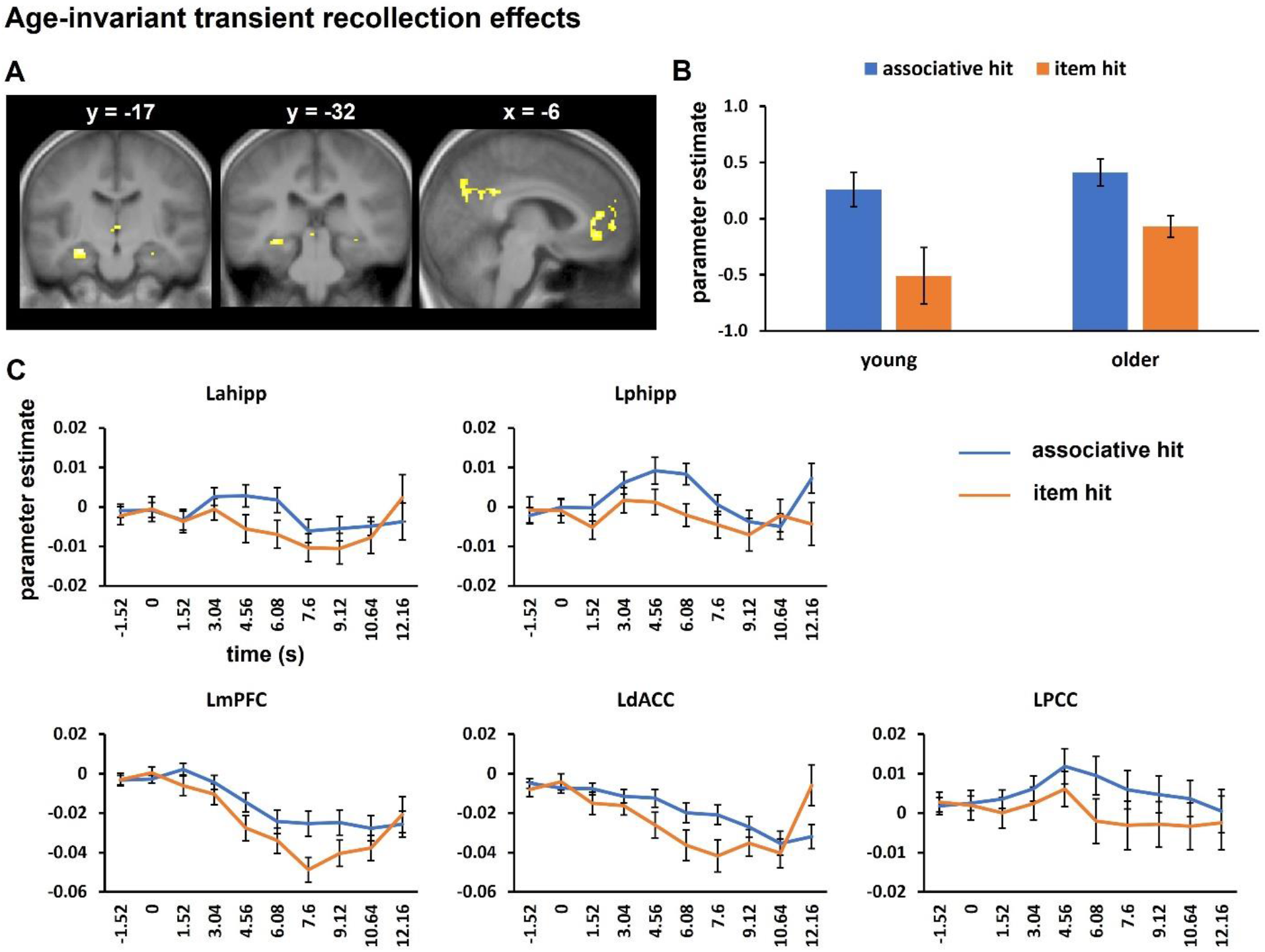
A: Clusters demonstrating age-invariant transient recollection effects. B: Parameter estimates derived from associative hit and item hit trials averaged across all of the regions illustrated in A. C: Time course data extracted from each region. In both B and C, and in succeeding figures, error bars indicate standard errors of the mean.

**Figure 3.**
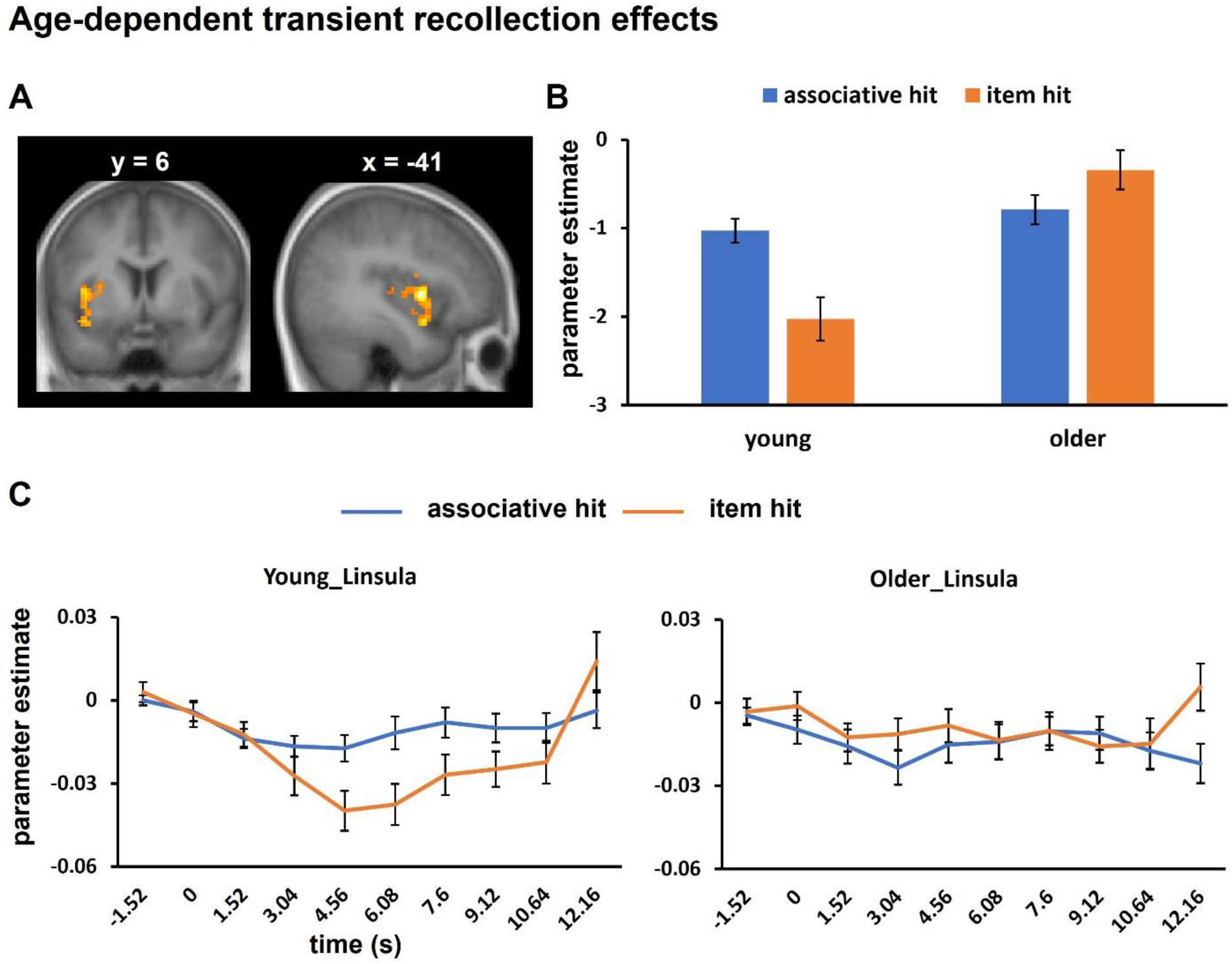
A: Cluster in left insula demonstrating a group difference in transient recollection effects. B: Mean parameter estimates. C. Group-wise time course data.

**Figure 4.**
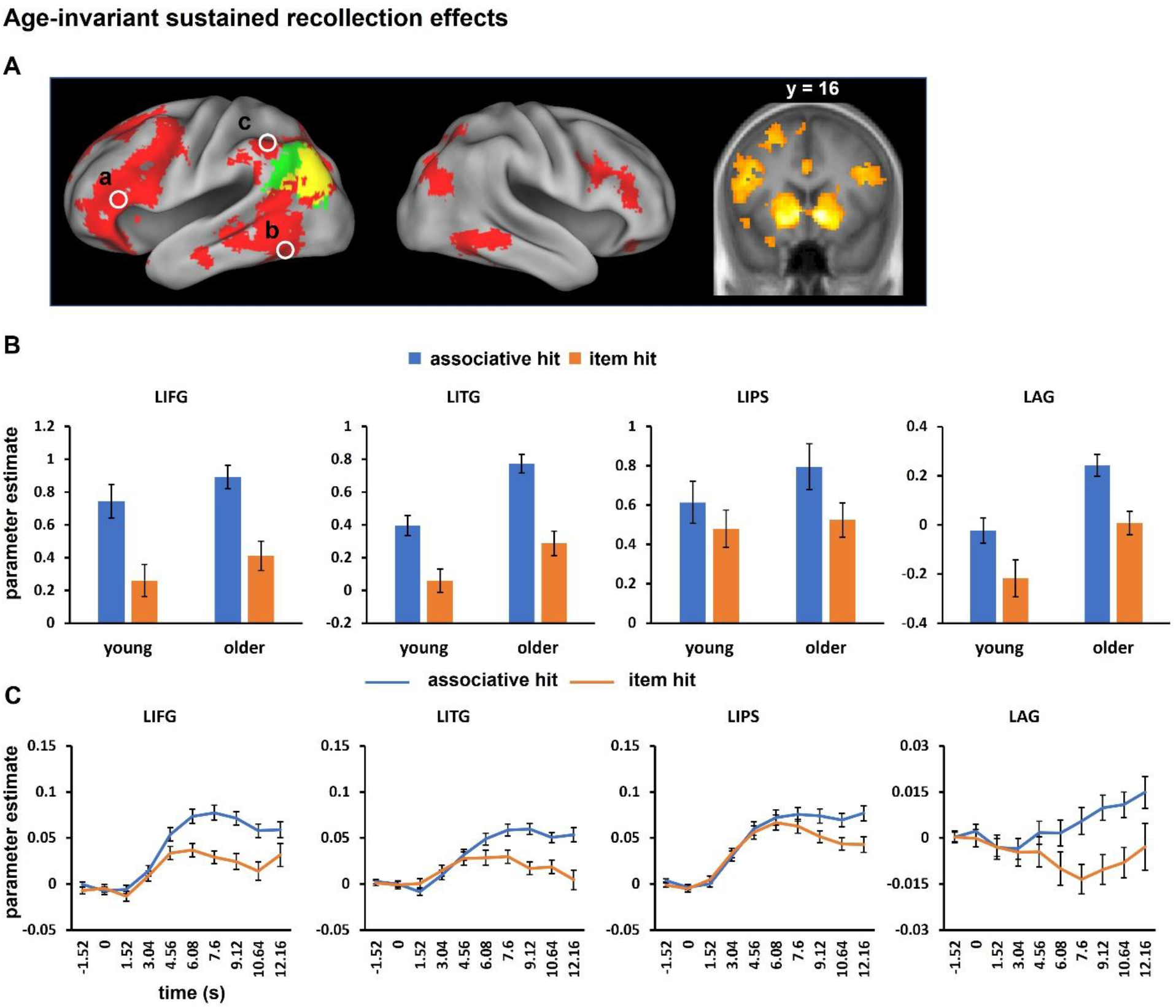
A: age-invariant sustained recollection effects. a. IFG (peak coordinate −48, 32, 13), b. ITG (−57, −46, −14), c. IPS (−33, −55, 46) and AG (ROI outlined in green and yellow - yellow demotes a spatial overlap between the ROI and the age-invariant recollection effect). B. Parameter estimates from the standard GLM. C: Time course data.

## 3 Results

### 3.1 Neuropsychological test performance

Demographic information and the neuropsychological test scores for the young and older age groups are summarized in Table 1. The sex distribution did not significantly differ between age groups, χ^2^ (1) = 0.57, p = 0.449. Older adults had more years of education than young adults, while young adults demonstrated higher performance on CVLT short delay free recall, SDMT, Trails A, Category Fluency and Raven’s Progressive Matrices. These findings are typical of those reported previously for similar samples of young and older adults (e.g., de Chastelaine et al., 2016).

**Table 1.** Demographic and neuropsychological data for young and older adults (standard deviations in parentheses).

|  | Young | Older |
| --- | --- | --- |
| N (sex) | 23 (11 F, 12 M) | 22 (15 F, 9 M) |
| Age | 21.87 (3.02) | 69.82 (3.22) |
| Years of education<sup>1</sup> | 15.37 (1.82) | 17.64 (2.17) |
| MMSE | 28.74 (1.18) | 28.95 (0.90) |
| CVLT short delay free recall<sup>1</sup> | 12.87 (2.38) | 10.68 (2.92) |
| CVLT short delay cued recall | 13.17 (2.21) | 12.50 (2.28) |
| CVLT long delay free recall | 12.83 (2.44) | 11.23 (2.88) |
| CVLT long delay cued recall | 13.13 (2.28) | 12.55 (2.60) |
| CVLT recall composite | 13.00 (2.13) | 11.74 (2.49) |
| CVLT recognition hits | 15.17 (0.98) | 15.23 (1.11) |
| CVLT recognition false alarms | 1.09 (1.38) | 2.18 (2.36) |
| Logical memory I | 30.22 (7.19) | 30.05 (6.78) |
| Logical memory II | 28.13 (6.51) | 27.18 (7.16) |
| Logical memory composite | 29.17 (6.68) | 28.61 (6.85) |
| F-A-S | 43.35 (13.11) | 47.09 (10.64) |
| SDMT<sup>1</sup> | 61.43 (12.31) | 49.77 (8.88) |
| Trails A<sup>1</sup> | 21.03 (7.39) | 29.90 (14.45) |
| Trails B | 52.22 (32.77) | 57.24 (19.58) |
| Digit span | 19.00 (5.25) | 18.50 (4.51) |
| Category fluency<sup>1</sup> | 25.91 (5.37) | 22.09 (3.37) |
| TOPF/WTAR est. full scale IQ | 109.43 (8.27) | 114.32 (10.27) |
| Raven's<sup>1</sup> | 11.09 (1.20) | 9.77 (1.95) |

Note. 1: young ≠ older, $p < 0.05$ .

### 3.2 Associative memory performance

Table 2 summarizes associative memory performance for the two age groups. A 2 (age group) x 3 (judgment type: associative hit, associative miss, correct rejection) ANOVA conducted on the response proportions gave rise to a significant main effect of judgment type (F_1.58, 68.05_ = 158.26, p < 0.001, partial η^2^ = 0.79) and a judgment type x age group interaction (F_1.58, 68.05_ = 5.92, p = 0.008, partial η^2^ = 0.12). The main effect of age group was not significant (F_1, 43_ = 2.92, p = 0.094, partial η^2^ = 0.06). Follow-up independent sample t-tests revealed significantly lower associative hit rates (t_37.52_ = 2.81, p = 0.008, Cohen’s d = 0.83) and higher associative miss rates (t_43_ = 2.41, p = 0.020, Cohen’s d = 0.72) in the older adults; no age group difference was identified for correct rejection rates (t_43_ = 1.04, p = 0.304, Cohen’s d = 0.31). No significant age group difference was identified for item memory (Pr), t_43_ = 1.57, p = 0.123, Cohen’s d = 0.47.

**Table 2.** Associative memory performance for young and older adults (standard deviations in parentheses).

|  | Proportions |  | RTs |  |
| --- | --- | --- | --- | --- |
|  | Young | Older | Young | Older |
| *Old items* |  |  |  |  |
| **Associative hit** | **0.51 (0.19)** | **0.38 (0.12)** | **1268 (303)** | **1694 (380)** |
| **Associative miss** | **0.35 (0.14)** | **0.43 (0.08)** | **1265 (358)** | **1549 (466)** |
| Yes/No incorrect | 0.11 (0.05) | 0.15 (0.09) | 1493 (463) | 1937 (514) |
| Old don't remember | 0.24 (0.12) | 0.28 (0.12) | 1182 (364) | 1325 (531) |
| Item miss | 0.13 (0.10) | 0.17 (0.11) | *1136 (472)* | 1044 (431) |
| **Pr** | **0.80 (0.15)** | **0.73 (0.12)** |  |  |
| *New items* |  |  |  |  |
| False alarm | 0.06 (0.10) | 0.08 (0.08) | 1148 (491) | 1727 (565) |
| Yes/No incorrect | 0.01 (0.03) | 0.04 (0.06) | *1562 (1476)* | *1902 (573)* |
| Old don't remember | 0.05 (0.09) | 0.04 (0.04) | *1144 (368)* | *1374 (680)* |
| **Correct rejection** | **0.92 (0.11)** | **0.88 (0.14)** | **820 (296)** | **879 (290)** |

Note. Statistics with missing data points are presented in italics. Data presented in bold was subjected to statistical analysis.

Reaction times (RTs), measured with respect to the onset of the post-delay cue, are also given in Table 2. A 2 (age group: young, older) x 3 (judgment type: associative hit, associative miss, correct rejections) ANOVA revealed significant main effects of judgment type (F_1.76, 75.53_ = 147.97, p < 0.001, partial η^2^ = 0.78) and age group (F_1, 43_ = 7.36, p = 0.010, partial η^2^ = 0.15), which were modified by a judgment type x age group interaction (F_1.76, 75.53_ = 10.59, p < 0.001, partial η^2^ = 0.20). Follow-up analyses revealed significantly longer RTs in the older adults for associative hits (t_43_ = 4.17, p < 0.001, Cohen’s d = 1.24) and misses (t_43_ = 2.30, p = 0.026, Cohen’s d = 0.69), but not for correct rejections, t_43_ = 0.69, p = 0. 497, Cohen’s d = 0.20.

We also contrasted associative hit rates according to age group and delay interval. The ANOVA revealed higher associative hit rates in young adults than older adults for all intervals. Of importance, the age group x interval interaction was non-significant (p = 0.382; see supplementary material).

### 3.3 fMRI results

#### 3.3.1 Age-invariant transient recollection effects

Five clusters were identified that demonstrated an age-invariant transient recollection effect. As is evident from Figure 2A and Table 4, the clusters were located in left anterior and posterior hippocampus, mPFC, dorsal anterior cingulate (dACC) and PCC. Mean parameter estimates averaged across these regions are shown in Figure 2B, while Figure 2C illustrates the time course data from each region (see supplemental Figure 1 for the time-courses for each age group).

**Table 4.**
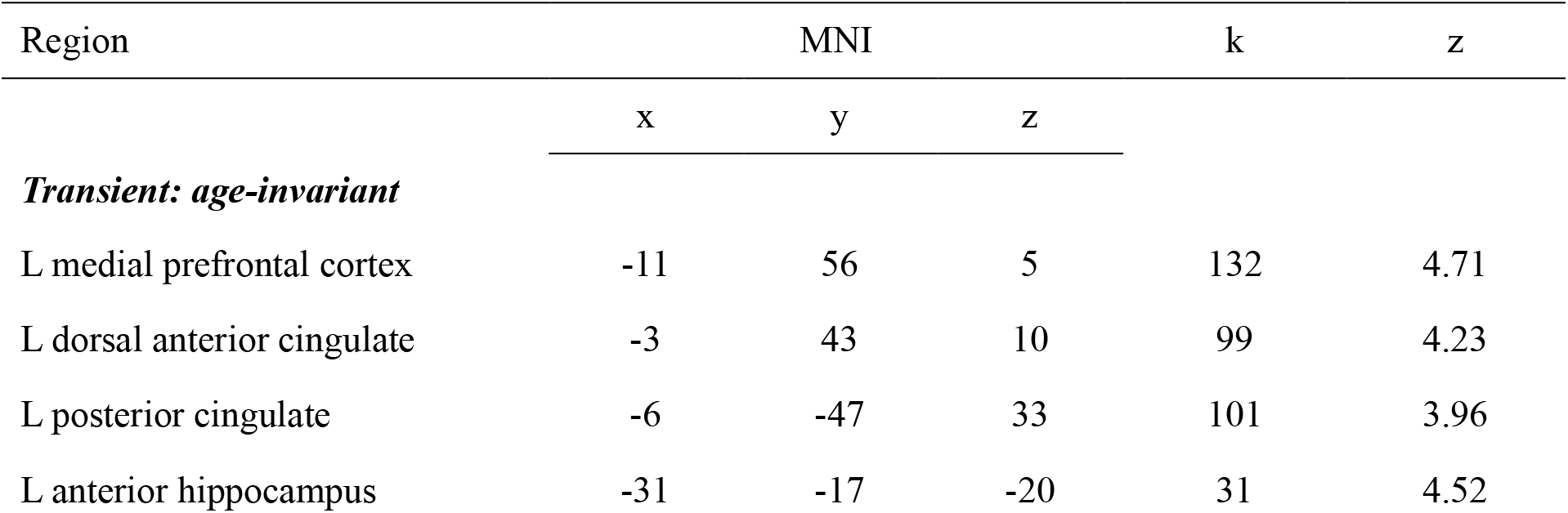
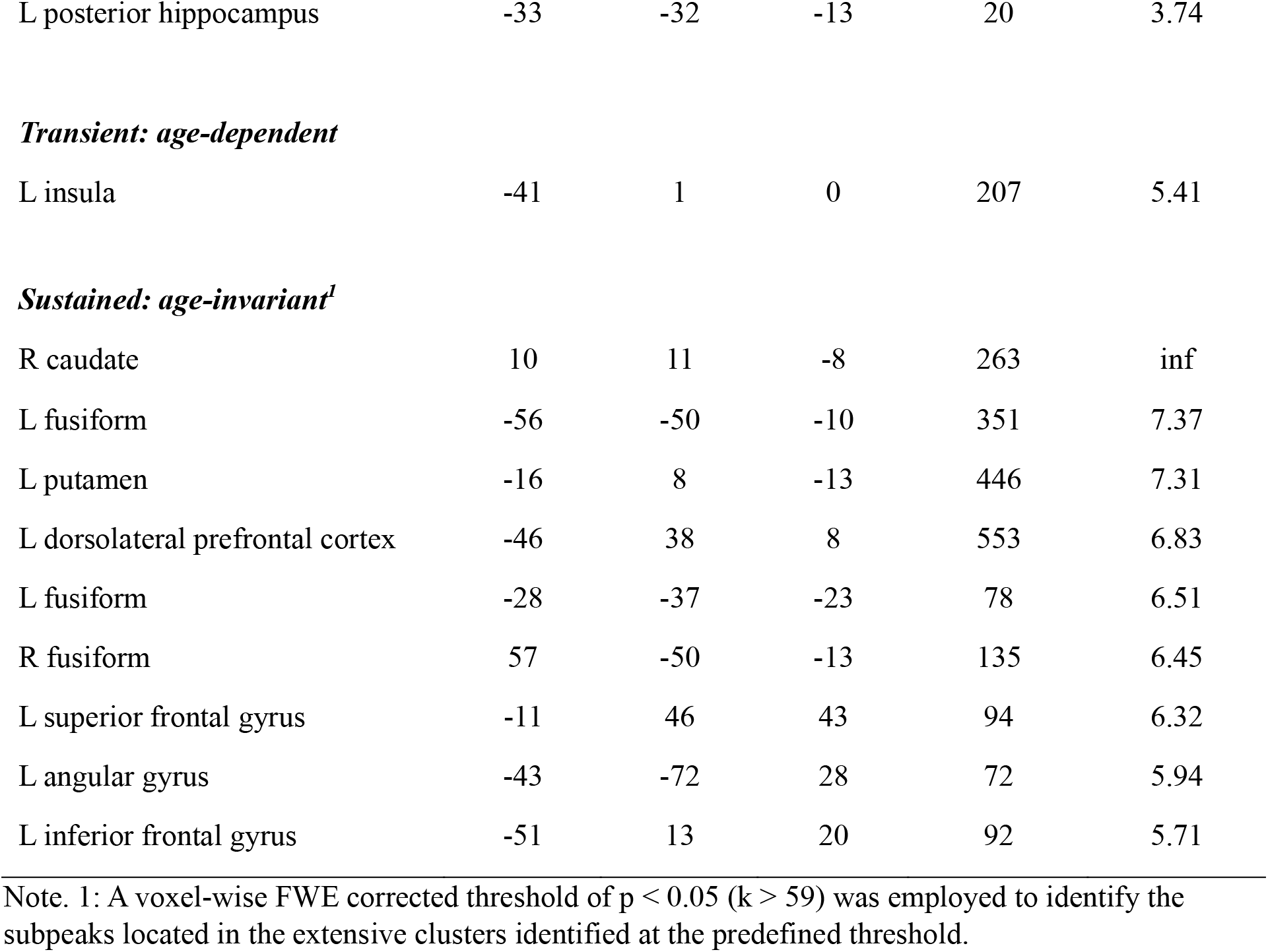
Regions demonstrating transient and sustained recollection effects identified from the whole brain analysis. The MNI coordinate of the peak of each cluster is listed.

#### 3.3.2 Age-dependent transient recollection effects

At our pre-defined threshold (see Methods), the age group x judgment type (associative hit, item hit) interaction contrast identified a single cluster in the left anterior insula (see Figure 3 and Table 4). A 2 (age group) x 2 (judgment type) ANOVA conducted on the mean parameter estimates from this region revealed a main effect of age group (F_1, 43_ = 16.25, p < 0.001, partial η^2^ = 0.27) and, of necessity, an age group x judgment type interaction (F_1, 43_ = 24.93, p < 0.001, partial η^2^ = 0.37). Follow-up pairwise t-tests revealed a significant recollection effect in the young group (t_22_ = 5.80, p < 0.001, Cohen’s d = 1.21) but a nonsignificant effect in the older adults (t_21_ = 1.91, p= 0.070, Cohen’s d = 0.41, see Figure 3B).

#### 3.3.3 Sustained recollection effects

No above-threshold clusters were identified by the age group x judgment type interaction contrast. By contrast, age-invariant effects were identified in multiple brain regions, including bilateral PFC, MTG, ITG, superior parietal cortex, AG, striatum, PCC and right cerebellum (see Figure 4A and Table 4).

As described in the Methods, time course data was extracted from four exemplary left cortical regions. These data are plotted in Figure 4C (see supplemental Figure 2 for the time-courses for each age group). Following Vilberg and Rugg (2012), we went on to examine whether recollection-related activity in the regions identified above tracked the delay interval. For each region we conducted a 2 (age group) x 2 (delay interval: 2s, 8s) x 11 (time point: 0-10 TRs after word onset) ANOVA on the parameter estimates for the associative hit trials derived from the FIR model. Table 5 shows the outcomes of the ANOVAs. As is evident from the table, the delay interval x time point interaction was reliable in each region. We followed up these interactions with pairwise t-tests contrasting the parameter estimates according to time point. As can be seen in Figure 5, the parameter estimates for the final two time points were significantly greater for the 8s than for the 2s delay in all regions (see supplemental material for full results).

**Table 5.** Results of ANOVAs comparing associative hits for the 2s vs. 8s delay interval in young and older adults.

| Source | df | df error | F | p | Partial $\eta^2$ |
| --- | --- | --- | --- | --- | --- |
| *LIFG* |  |  |  |  |  |
| Delay interval | 1 | 44 | 2.23 | 0.142 | 0.05 |
| **Time point** | **4.02** | **176.85** | **40.59** | **< 0.001** | **0.48** |
| **Delay interval x time point** | **4.79** | **210.87** | **11.09** | **< 0.001** | **0.20** |
| *LITG* |  |  |  |  |  |
| **Delay interval** | **1** | **44** | **8.00** | **0.007** | **0.15** |
| **Time point** | **3.64** | **159.94** | **27.91** | **< 0.001** | **0.39** |
| **Delay interval x time point** | **4.74** | **208.59** | **7.69** | **< 0.001** | **0.15** |
| *LIPS* |  |  |  |  |  |
| **Delay interval** | **1** | **44** | **12.44** | **< 0.001** | **0.22** |
| **Time point** | **3.37** | **148.48** | **44.45** | **< 0.001** | **0.50** |
| **Delay interval x time point** | **4.42** | **194.51** | **29.80** | **< 0.001** | **0.40** |
| *LAG* |  |  |  |  |  |
| Delay interval | 1 | 44 | 2.72 | 0.106 | 0.06 |
| **Time point** | **3.61** | **158.69** | **4.99** | **0.001** | **0.10** |
| **Delay interval x time point** | **3.72** | **163.66** | **3.80** | **0.007** | **0.08** |

**Figure 5.**
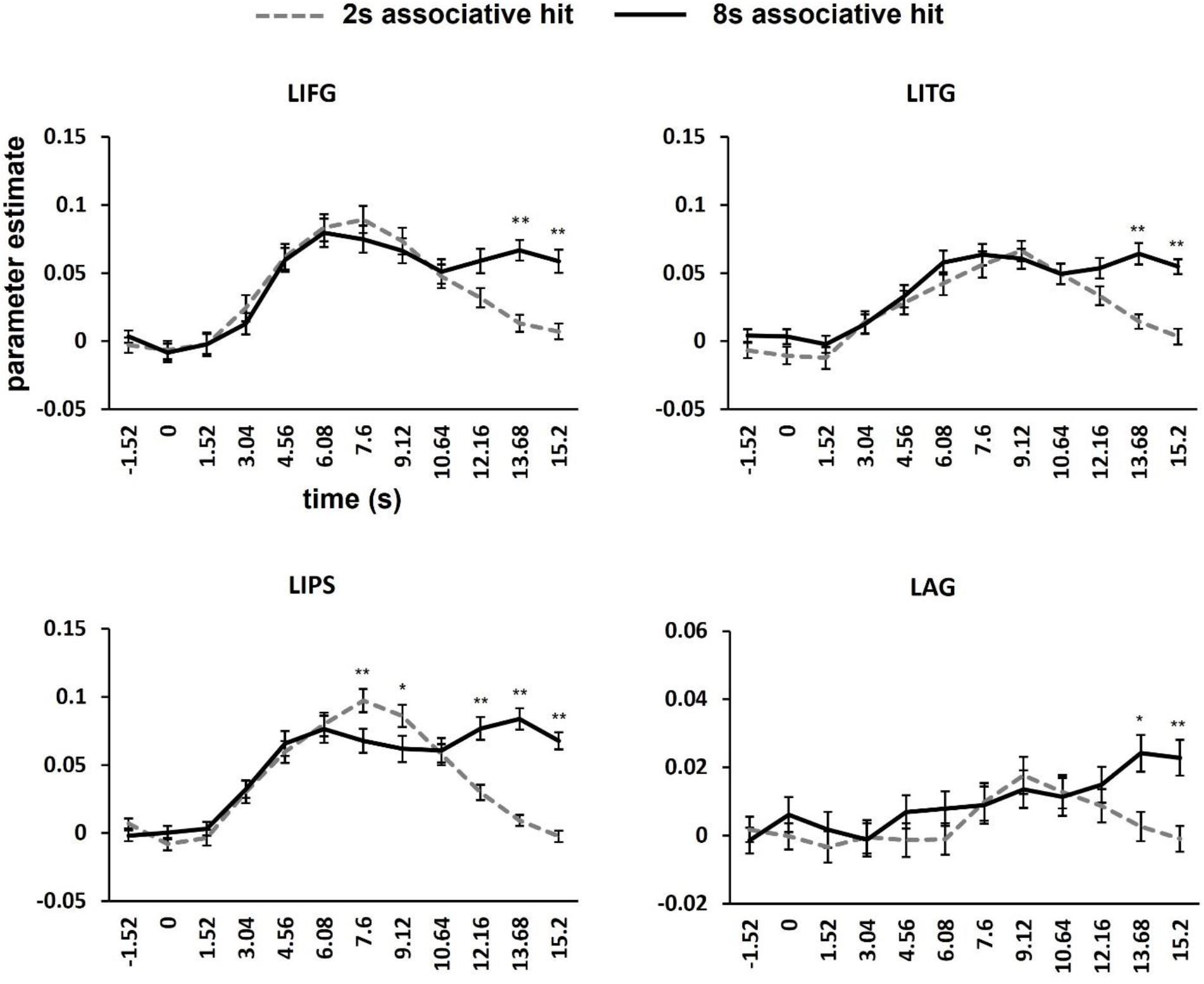
FIR parameter estimates from left IFG, ITG, IPS and AG are displayed separately for the 2 and 8 s delay interval associative hit trials. Significant findings in each region that survived correction for multiple comparisons (p = 0.0045) are indicated. *p < 0.0045, **p < 0.002.

## 4. Discussion

The present study examined transient and sustained fMRI recollection effects in young and older adults. Replicating the findings of Vilberg and Rugg (2012, 2014), transient effects were identified in the left hippocampus, mPFC and PCC, while sustained effects were widespread across the left lateral cortex and were also evident in the bilateral striatum. Remarkably, with the exception of the left insula, these effects, whether transient or sustained, were age-invariant. These findings extend prior reports by suggesting that recollection effects, both within and outside of the core recollection network, are largely age-invariant even when the recollected information must be held in mind for almost 10s.

### 4.1 Behavioral results

As is often reported (for reviews, see Koen and Yonelinas, 2014; Old and Naveh-Benjamin, 2008), while item (recognition) memory did not significantly differ according to age, associative memory performance was lower in the older than the young age group. The magnitude of the age difference in memory performance did not significantly vary with delay interval, however. Thus, although older adults were disadvantaged relative to their younger counterparts in recollecting associative information in response to a retrieval cue, this disadvantage did not extend to the ability to maintain recollected information in working memory (on the assumption that a working memory impairment would have led to a negative association between accuracy and delay interval). At first sight, this finding conflicts with prior studies in which older adults were reported to demonstrate lower delayed response performance than young participants (e.g., Cappell et al., 2010; Benett et al., 2013). However, the memory loads in these prior studies were invariably higher than the load imposed here (2 or more items, rather than the single item in the current study). From this perspective, it would be expected that, had our older adults been required to retrieve and maintain multiple study items, accuracy would likely have declined with increasing delay.

### 4.2 Transient recollection effects

Turning to the fMRI findings, we identified age-invariant transient recollection effects in the left hippocampus, mPFC and PCC, along with an age difference in the left insula. The findings for the hippocampus and midline cortical regions are closely similar to those reported by Vilberg and Rugg (2012, 2014). As was discussed by those authors, the transient effects evident in the hippocampus are consistent with the idea that while this region is responsible for reinstating memory representations in the neocortex (e.g., Norman and O’Reilly, 2003; Rugg et al., 2008; Staresina and Wimber, 2019), it plays little or no role in subsequently maintaining these representations. Relatedly, the transient effects evident in PCC and mPFC might also reflect a role, in concert with the hippocampus, in accessing stored episodic information, perhaps facilitating access through the retrieval of gist- or schema-based information (Baldassano et al., 2018; Ranganath and Ritchey, 2012; Reagh and Ranganath, 2023; Ritchey and Cooper, 2020). Regardless of their precise functional significance, a key finding is that as we had expected, the transient effects identified in these medial temporal and midline core recollection regions were age-invariant. This finding adds to the evidence that the neural correlates of recollection in these regions are largely insensitive to increasing age (see Introduction).

Unexpectedly, we identified an effect of age on transient recollection effects in the left insula, where effects were highly robust in the young age group, but absent in the older adults. Although it is rarely mentioned as a recollection-sensitive neural region, reports of recollection effects in this part of the insula are not unprecedented (Kim, 2020), although their functional significance is obscure. Motivated by the present finding, we conducted a retrospective analysis of data acquired in two earlier studies examining the effects of age on recollection effects that were conducted in the authors’ laboratory (de Chastelaine et al., 2016; Wang et al, 2016). In both cases, we identified robust recollection effects in the same left insula region as was identified in the present study. However, in contrast to the present findings, there was no evidence of an effect of age in these two datasets. In light of these null findings, we refrain from offering an interpretation of the present finding until it has been demonstrated to be reproducible.

### 4.3 Sustained recollection effects

The primary goal of the present study was to determine whether sustained recollection effects are age-sensitive. Reassuringly, the sustained effects identified in the young age group took the form of a full replication of those reported previously (Vilberg and Rugg, 2012, 2014). Crucially, these effects were equally robust in our older participants and, so far as we could tell, did not significantly differ from those of the young group either in their magnitudes or their regional extents. Thus, the present findings demonstrate an age-invariant dissociation between the temporal properties of recollection effects in midline and medial temporal (PCC, mPFC, hippocampus) and lateral cortical (left AG and MTG) members of the core recollection network. Although the functional significance of MTG recollection effects remains unclear, the left AG has long been proposed to serve as some sort of ‘episodic buffer’ capable of representing and integrating multi-featural episodic information (Guerin and Miller, 2011; Humphreys et al., 2021; Rugg and King, 2018; Vilberg and Rugg, 2008). The present findings add weight to this proposal and suggest that any such buffering function is preserved in cognitively healthy older adults.

Sustained recollection effects were prominent not only in the MTG and AG, but in lateral prefrontal and dorsal posterior parietal cortex, as well as in the striatum. These regions, which are held to comprise a functional network underpinning cognitive control (the ‘fronto-parietal control network’), have frequently been reported to be active during the delay period of delayed response tasks (for review, see D’Esposito and Postle, 2015). It has been proposed that the network implements control processes that, among other functions, support the maintenance of information in working memory (e.g., Jonides et al., 2008; Wager and Smith, 2003; but see Riggall and Postle, 2012, and Postle, 2015). Given that the present task required the maintenance of task-relevant information across a delay, it is unsurprising that similar control processes were engaged during its performance.

The present study has several limitations. First, the sample sizes are relatively small, signaling the need for caution in accepting null findings for the effects of age (although any such effects, should they exist, seem likely to be modest). Second, we employed a cross-sectional design and therefore cannot distinguish between the influences of age and age-related confounds such as cohort effects or selection bias (Rugg, 2017). Third, the fMRI BOLD signals we employed to estimate neural activity might have been confounded with age differences in vascular factors that mediate between neural activity and the BOLD signal (e.g., Liu et al. 2013, Lu et al. 2011, Tsvetanov et al. 2015). Fourth, although we identified regional dissociations in the temporal properties of recollection effects, we were unable to precisely characterize their time courses due to the low temporal resolution of BOLD fMRI. Future research employing methods with higher temporal resolution (e.g., electrophysiological methods) would be valuable in advancing this line of research and, indeed, might reveal age differences in the timing of recollection effects that are undetectable with fMRI.

## 5. Conclusions

The present study adds to the evidence that recollection effects in different neural regions are temporally dissociable. Of more importance in the present context, the findings suggest that both transient and sustained recollection effects are largely insensitive to the effects of advancing age. Thus, the findings add significant and novel support to the contention that the neural correlates of recollection and, by extension, the functional integrity of recollection-sensitive neural regions, varies little across much of the adult lifespan.

## Supporting information

Hou_supplementary

## Acknowledgments

This work was supported by the National Institute on Aging (grant number RF1AG039103). We thank Amber Kidwai, Ayse Aktas, Chris Hawkins, Eduardo Hernandez, Joshua Oliver, Melanie Racenstein and Seham Kafafi, for their contributions to the recruitment and neuropsychological testing of the experimental participants.

