## Supplementary material for "The effects of age on neural correlates of recollection: transient versus sustained fMRI effects": Hou_supplementary

1. Comparisons of associative hit rates across delay intervals

2. Supplemental Figure 1. Time course data from each region demonstrating age-invariant transient recollection effects in each age group.

3. Supplemental Figure 2. Time course data from selected regions demonstrating age-invariant sustained recollection effects in each age group.

4. Supplemental Figure 3. FIR parameter estimates for left IFG, ITG, IPS and AG for the 2 and 8 s delay interval associative hit trials shown separately for each age group.

5. Supplemental Table 2. Results of pairwise t-tests comparing parameter estimates for associative hits from the 2s vs. 8s delay intervals.

6. Supplemental Table 3. Regions demonstrating transient and sustained recollection effects identified from whole brain analyses employing a 10-trial cutoff. The MNI coordinates of the peak of each cluster are listed.

Comparisons of associative hit rates across delay intervals

Associative hit rates for each delay interval are given in Supplemental Table 1. The data were analyzed with a 4 (delay interval: 2, 4, 6, 8s) x 2 (age group) ANOVA. The ANOVA gave rise to a significant effect of age group (F_1, 43_ = 7.72, p = 0.008, partial ƞ^2^ = 0.15. Neither the main effect of delay interval or the delay interval x age group interaction was significant; respectively, F_3, 129_ = 0.55, p = 0.647, partial ƞ^2^ = 0.01 and F_3, 129_ = 1.03, p = 0.382, partial ƞ^2^ = 0.02).

Supplemental Table 1. Associative hit rates according to age group and delay interval (standard deviations in parentheses).

|  | Delay interval |  |  |  |
| --- | --- | --- | --- | --- |
|  | 2s | 4s | 6s | 8s |
| Young | 0.34 (0.14) | 0.31 (0.12) | 0.32 (0.14) | 0.31 (0.12) |
| Older | 0.23 (0.10) | 0.24 (0.08) | 0.25 (0.11) | 0.23 (0.09) |

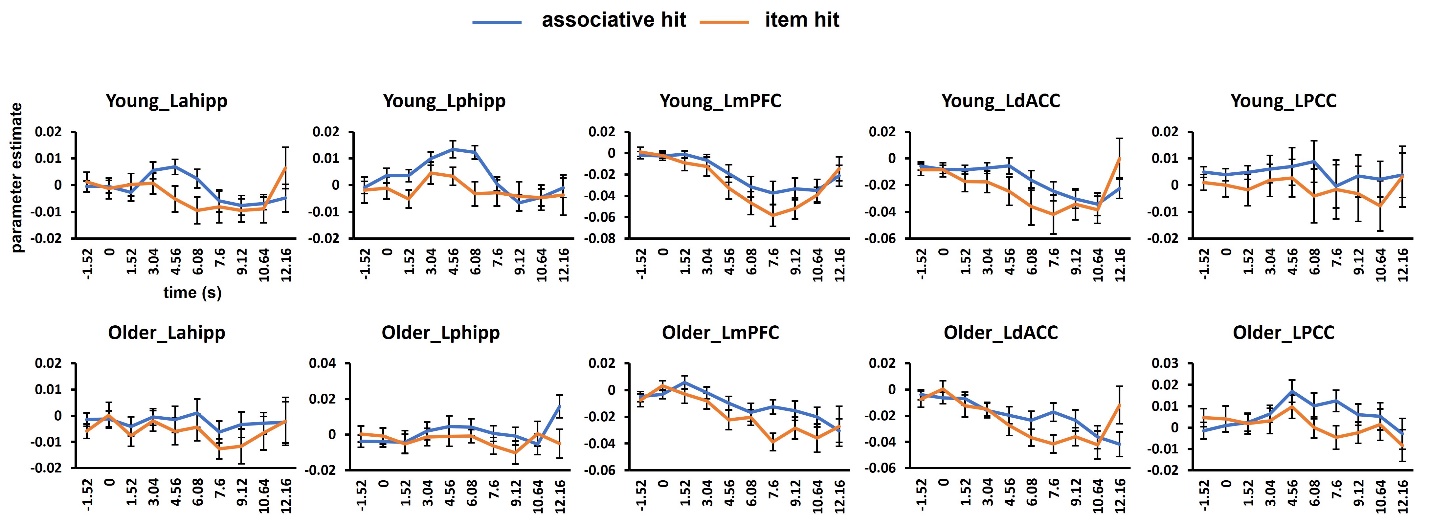

Supplemental Figure 1. Time course data from each region demonstrating transient recollection effects in each age group.

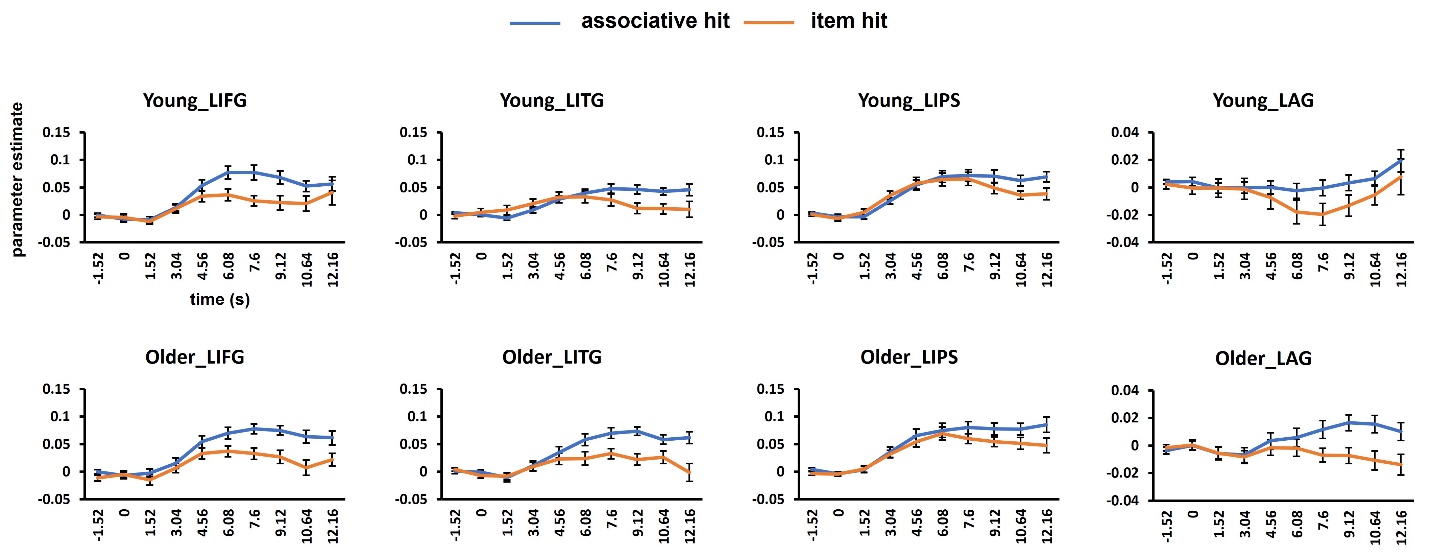

Supplemental Figure 2. Time course data from selected regions demonstrating sustained recollection effects in each age group.

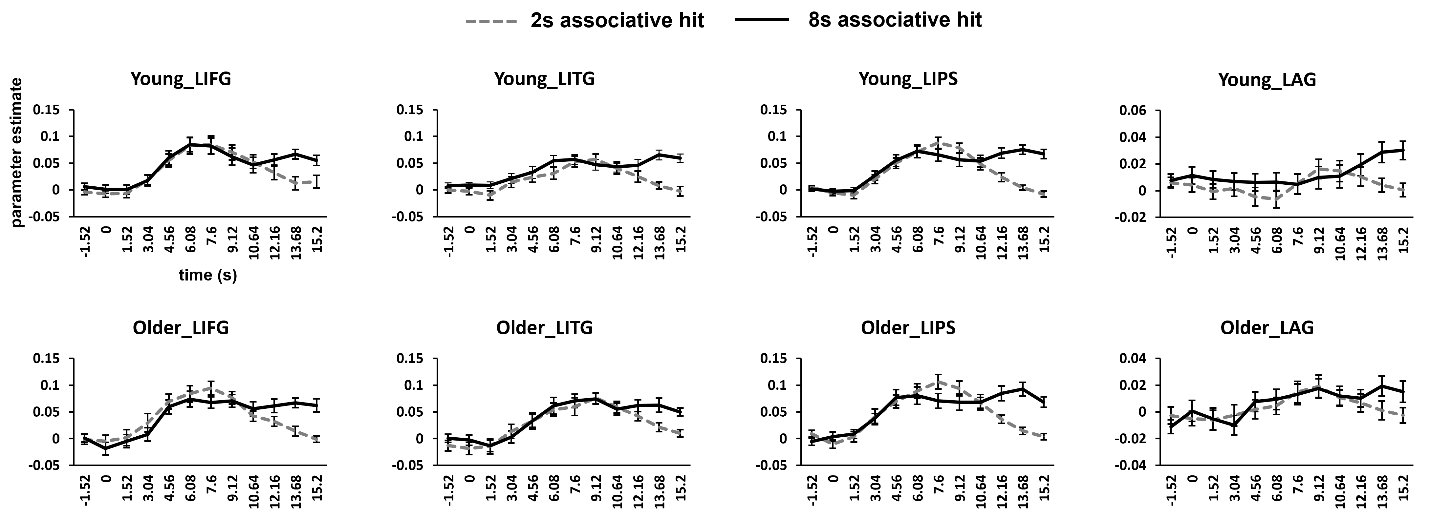

Supplemental Figure 3. FIR parameter estimates for left IFG, ITG, IPS and AG for the 2 and 8 s delay interval associative hit trials shown separately for each age group.

Supplemental Table 2. Results of pairwise t-tests comparing parameter estimates for associative hits from the 2s vs. 8s delay intervals.

|  | LIFG | LITG | LIPS | LAG |
| --- | --- | --- | --- | --- |
| 0s | p = 0.813 | p = 0.119 | p = 0.246 | p = 0.313 |
| 1.52s | p = 0.988 | p = 0.265 | p = 0.387 | p = 0.425 |
| 3.04s | p = 0.257 | p = 0.859 | p = 0.812 | p = 0.878 |
| 4.56s | p = 0.781 | p = 0.569 | p = 0.261 | p = 0.108 |
| 6.08s | p = 0.668 | p = 0.060 | p = 0.601 | p = 0.069 |
| 7.6s | p = 0.103 | p = 0.399 | **2s > 8s, p < 0.001** | p = 0.866 |
| 9.12s | p = 0.346 | p = 0.432 | **2s > 8s, p = 0.002** | p = 0.504 |
| 10.64s | p = 0.710 | p = 0.998 | p = 0.689 | p = 0.809 |
| 12.16s | 2s < 8s, p = 0.005 | 2s < 8s, p = 0.031 | **2s < 8s, p < 0.001** | p = 0.314 |
| 13.68s | **2s < 8s, p < 0.001** | **2s < 8s, p < 0.001** | **2s < 8s, p < 0.001** | **2s < 8s, p = 0.002** |
| 15.2s | **2s < 8s, p < 0.001** | **2s < 8s, p < 0.001** | **2s < 8s, p < 0.001** | **2s < 8s, p = 0.001** |

Note. Results that survived correction for multiple comparisons (threshold p = 0.0045) are shown in bold.

Supplemental Table 3. Regions demonstrating transient and sustained recollection effects identified from whole brain analyses employing a 10-trial cutoff (data from 20 young, and 20 older participants). The MNI coordinate of the peak of each cluster is listed.

| Region | MNI | | | k | z |
| --- | --- | --- | --- | --- | --- |
|  | x | y | z |  |  |
| ***Transient: age-invariant*** |  |  |  |  |  |
| R medial prefrontal cortex | 2 | 41 | 10 | 244 | 4.63 |
| L precuneus | -6 | -72 | 30 | 81 | 4.06 |
| L anterior hippocampus | -31 | -17 | -20 | 18 | 4.26 |
| L posterior hippocampus | -33 | -32 | -13 | 15 | 3.83 |
| R anterior hippocampus | 30 | -20 | -20 | 11 | 3.58 |
| ***Transient: age-dependent*** |  |  |  |  |  |
| L insula | -41 | 1 | 0 | 152 | 5.18 |
| ***Sustained: age-invariant^1^*** |  |  |  |  |  |
| R caudate | 10 | 8 | -8 | 386 | inf |
| L fusiform | -56 | -50 | -10 | 341 | 7.80 |
| L putamen | -16 | 6 | -15 | 476 | 7.70 |
| L dorsolateral prefrontal cortex | -46 | 38 | 8 | 703 | 6.93 |
| L fusiform | -28 | -37 | -23 | 66 | 6.27 |
| R fusiform | 57 | -50 | -13 | 99 | 6.23 |
| L superior frontal gyrus | -11 | 46 | 43 | 82 | 6.16 |
| L angular gyrus | -43 | -72 | 28 | 132 | 5.96 |
| L inferior/middle frontal gyrus | -43 | 13 | 33 | 261 | 5.94 |
| L middle frontal gyrus | -28 | 21 | 50 | 60 | 5.79 |

Note. 1: A voxel-wise FWE corrected threshold of p < 0.05 (k > 59) was employed to identify the subpeaks located in the extensive clusters identified at the predefined threshold.
